# Structural Variant Aware Identity-by-Descent Detection on Pangenome Graphs

**DOI:** 10.64898/2026.11.05.764670

**Authors:** Uwaise Islam, Franco Marsico, Erik Garrison, Christina Boucher

## Abstract

Identity-by-descent (IBD) detection identifies genomic segments inherited from a common ancestor, and supports studies of relatedness, demographic history, and disease mapping. Existing methods primarily use linear genome representations and do not identify structural variants (SVs) shared within IBD segments. Pangenome graphs encode SVs and store each haplotype as a path through the graph. We present GIBD (Graph Identity-by-Descent), which detects IBD segments in the Graph Burrows–Wheeler Transform (GBWT) of a pangenome graph and annotates each detected segment with the SVs. GIBD enumerates the pairwise-maximal exact matches between all pairs of haplotype paths in a single traversal of the GBWT, and visits each shared subpath once. On graphs of bounded degree, the traversal takes time linear in the number of nodes on the haplotype paths and the number of matches it enumerates. On a standard IBD benchmark, GIBD’s power and accuracy are within 0.009 and 0.001 of `hap-ibd`, a widely used linear method. Compared with `impop`_*k*_, a pangenome-based IBD method, GIBD detects more 2–4 cM segments and runs in 33 seconds instead of 9.6 hours. In simulations of 1,000 haplotypes built from assemblies of the Human Pangenome Reference Consortium (HPRC), GIBD’s insertion, deletion, and inversion labels have a precision of 1.000, with and without assembly error. In a real four-generation family, GIBD identifies the inherited allele on parent–offspring segments at all 3,235 SVs on chromosome 20 where the parent is heterozygous, and reports no IBD between unrelated members. It also tracks a disease-associated 900-kb inversion at 17q21.31 across three generations and correctly labels all segments that share it. GIBD makes SVs visible in IBD detection from pangenome graphs.

## 1 Introduction

Identity-by-descent (IBD) segments are genomic regions inherited from a common ancestor without intervening recombination. Detecting these segments is a fundamental task in population and medical genomics, supporting analyses of relatedness, demographic history, and the inheritance of disease-associated variation [1, 2]. Two haplotypes that share an IBD segment carry the same alleles throughout the segment, apart from mutations that arose after the common ancestor. These alleles include structural variants (SVs) such as insertions, deletions, inversions, and other changes of at least 50 bases [3]. SVs have been shown to affect disease and other traits [4, 5, 6, 7], yet their inheritance and distribution across populations remain difficult to characterize [8, 9]. An IBD detection method that identifies shared SVs within IBD segments would trace SV inheritance through families, characterize how SVs are distributed across populations, and help reveal their evolutionary origins. Linking SVs to shared ancestral haplotypes would also support disease-gene mapping and the interpretation of rare SVs [1, 10, 11, 3].

Existing IBD detection methods operate on haplotypes represented as sequences of alleles at ordered variant sites, mostly single nucleotide polymorphisms (SNPs), along a linear reference genome [12, 13, 14, 15]. We call these methods linear IBD methods. Ordered variant sites allow efficient matching across large haplotype collections [16], but this representation requires the same sites in the same order on every haplotype. SVs of different types break this assumption. An insertion adds a sequence that has no place on the reference, a deletion leaves only missing sites, and an inversion reverses a region whose sites still appear in reference order, so a linear method sees none of the three as an SV.

Pangenome graphs combine hundreds of genome assemblies, each the full DNA sequence of one haplotype, and natively represent SVs [9, 8]. Each haplotype is a path that walks through the nodes of a pangenome graph in the order of its sequence [17]. Where two haplotypes agree, their paths run through the same nodes in the same direction. Where they differ, their paths split. The Graph Burrows–Wheeler Transform (GBWT) stores all haplotype paths in compressed form and supports efficient matching between them [18]. A GBWT querying method, `GBWT-Query`, finds long matches between one query haplotype and all the others, so comparing all pairs takes one run per haplotype [19]. `impop`_*k*_ takes a complementary approach and infers IBD from the sequence identity of two haplotypes in windows of a reference genome [20]. No existing method enumerates the longest (pairwise-maximal) matches between any two haplotypes, where their paths run through exactly the same nodes, and joins them into IBD segments.

We present GIBD, to our knowledge, the first method that finds IBD segments in a pangenome graph and labels each segment with the SVs. GIBD enumerates the pairwise-maximal matches between all pairs of haplotypes in the GBWT, and then joins them into IBD segments [12]. Our main algorithmic contribution is a single traversal of the GBWT that visits each subpath that several haplotypes share only once, no matter how many haplotypes share it, much as a suffix tree lists the repeated substrings of a text [21, 22]. If every node has a bounded number of neighbors, GIBD enumerates these matches in *O*(*n* + *occ*) time, where *n* is the total number of nodes on all haplotype paths in both orientations and *occ* is the number of matching pairs of occurrences, which can grow with the square of the number of haplotypes.

GIBD labels each IBD segment with the insertions, deletions, and inversions that both haplotypes carry. Linear methods still report IBD segments that contain SVs, because the flanking SNPs match. This includes an inversion that only one of the two haplotypes carries. Many inversions arose repeatedly in human history, so an inversion can arise again on one of two haplotypes that share a common ancestor [23, 24]. GIBD reads SVs from the haplotype paths of the pangenome graph, so it can also label a segment in which only one haplotype carries an inversion.

GIBD’s pairwise-maximal matches are identical to the `GBWT-Query` output on 5,008 haplotypes of the 1000 Genomes Project [25, 19]. On the benchmark of linear IBD methods [26], GIBD matches the power and accuracy of `hap-ibd` at every genotyping-error rate, while `RaPID` and `TPBWT` reach full power on short segments at lower accuracy [12, 13, 15]. Compared with `impop`_*k*_ on simulated pangenome data, GIBD finds more short segments, with higher length accuracy [20]. We test GIBD’s SV labeling on haplotypes simulated from assemblies of the Human Pangenome Reference Consortium (HPRC) [27], so the IBD segments and SVs are known. On the segments GIBD detects, its insertion and deletion labels have precision and recall 1.000. When only one of the two haplotypes carries an inversion, GIBD labels every segment that spans it as differing by the inversion, with no false positive among all pairs it spans that do not differ. Since every offspring inherits one of the two haplotypes of each parent, the SVs where the parent is heterozygous show whether an IBD segment links the child to the right parental haplotype. In the Platinum Pedigree, a real four-generation family with assembled haplotypes [28], GIBD’s parent–offspring segments link each child to the right parental haplotype at all 3,235 such SVs on chromosome 20. The 900-kb inversion at 17q21.31 is a risk locus for progressive supranuclear palsy and the 17q21.31 microdeletion syndrome [29, 30]. In the same family, GIBD traces its H2 allele [7] across three generations and correctly labels all 6 segments that share it. Finally, GIBD’s time grows about linearly with the number of haplotypes and its memory more slowly, up to 462 haplotypes on chromosome 22 of the T2T-CHM13 HPRC graph.

## 2 Preliminaries

### Pangenome graph and GBWT

A pangenome graph represents a collection of genomes in a population as a bidirected graph *G* = (*V, E, P* ) [17]. Each node *v* ∈*V* is labeled with a genomic sequence *l*(*v*) ∈ {*A, C, G, T, N* } ^+^, and the edges in *E* connect nodes to encode sequence adjacencies and their relative orientations. The collection *P* consists of the paths corresponding to the input haplotypes and may not include all possible paths in *G*.

To support traversal of the haplotype paths *P* in compressed space, the Graph Burrows–Wheeler Transform (GBWT) stores these paths as a compressed and searchable index [18]. The GBWT is widely used in practice. For example, the human pangenome reference graph is distributed in a GBWT-based format that stores the GBWT together with the graph [31, 9].

To account for both DNA strand orientations and support BWT operations on bidirected graphs, we represent each node of *G* in both forward and reverse-complement orientations, yielding a graph *G*^*±*^ = (*V* ^*±*^, *E*^*±*^), where *V* ^*±*^ = *V*_*f*_ ∪ *V*_*r*_, where *V*_*f*_ and *V*_*r*_ contain the nodes in forward and reverse orientation, respectively. The sets *V*_*f*_ and *V*_*r*_ are disjoint, and |*V*_*f*_ | = |*V*_*r*_| = |*V* |. For each oriented node *v* ∈ *V* ^*±*^, 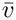 denotes the same underlying node in the opposite orientation. The operation satisfies 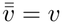. We extend *l* to oriented nodes so that 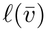 is the reverse complement of *l*(*v*). For all *u, v* ∈ *V* ^*±*^, the directed edge set satisfies 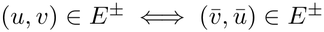. Each input path *P* = (*v*_1_, …, *v*_|*P* |_) ∈ *P* is indexed along with its reverse-complement path 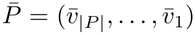. We denote this collection of indexed paths by *P*^*±*^, retaining a distinct identifier for each input path and each of its two orientations, even when their node sequences are identical.

To represent *P* ^*±*^, the GBWT stores a record BWT_*v*_ for each oriented node *v* ∈ *V* ^*±*^. Each record consists of a record body and a record header. The body of BWT_*v*_ contains a run-length encoded (RLE) sequence of successor nodes. Its rows represent visits to *v* along indexed paths, ordered colexicographically by the path prefixes ending at those visits, with ties broken by the path identifier. Each row stores the next node *w* ∈ *V* ^*±*^ ∪ {$}, where $ denotes path termination. We use zero-based row offsets throughout.

For each valid successor *w* ∈ *V* ^*±*^, the header of BWT_*v*_ stores a base offset *C*(*v, w*), the first row of BWT_*w*_ occupied by visits that arrive at *w* from *v*. If row *k* of BWT_*v*_ stores *w*, the target row is *k*^*′*^ = *C*(*v, w*) + rank_*w*_(BWT_*v*_, *k*), where rank_*w*_(BWT_*v*_, *k*) counts occurrences of *w* strictly before row *k*. Traversals are computed via the LF-mapping function LF(*v, k*) = (*w, k*^*′*^), which maps offset *k* at node *v* to offset *k*^*′*^ at successor node *w*. If the stored successor is $, the traversal terminates. The operations described here do not require an ordinary node record BWT_$_.

### Subpaths and occurrences

For an indexed path *P* = (*v*_1_, …, *v*_|*P* |_) ∈ *P*^*±*^, we denote *P* [*i*..*j*] = (*v*_*i*_, …, *v*_*j*_) (1 ≤ *i* ≤ *j* ≤ |*P* |) as the subpath from the *i*-th to the *j*-th node of *P* . It has |*P* [*i*..*j*]| = *j* − *i* + 1 nodes and a sequence length of 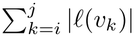 bases. Throughout, |*Q*| denotes the number of nodes in a subpath *Q*. All subpaths considered below are nonempty.

An *occurrence* of a subpath *Q* is a pair (*P, i*), where *P* ∈ *P*^*±*^ and *i* is the index of the node where *Q* starts, such that *P* [*i*..*j*] = *Q* with *j* = *i* + |*Q*| − 1 ≤ |*P* |. The set of all occurrences of *Q* in the indexed paths is denoted Occ(*Q*), with |Occ(*Q*)| representing the total number of occurrences. For an occurrence (*P, i*) of *Q* = *P* [*i*..*j*], the preceding node is *v*_*i−*1_ if *i >* 1, and the succeeding node is *v*_*j*+1_ if *j <* |*P*| . For boundary comparisons, we use $ to denote the absence of a preceding or succeeding node at a path boundary.

For a subpath *Q* ending at node *v*, all occurrences of *Q* correspond to a contiguous interval of rows [*s, e*] in BWT_*v*_, and |Occ(*Q*)| = *e* − *s* + 1. The node following an occurrence is the successor stored in its row. Because the rows are ordered colexicographically, occurrences preceded by a particular node *u* form a contiguous subinterval [*s*_*u*_, *e*_*u*_] ⊆ [*s, e*]. This subinterval represents exactly the occurrences of the longer subpath obtained by prepending *u* to *Q*. These subintervals are disjoint and, together with the rows representing occurrences that start at a path boundary, make up the interval [*s, e*]. However, the rows of [*s, e*] do not give [*s*_*u*_, *e*_*u*_]. Since the GBWT indexes all haplotype paths in both orientations, the subpath *Q* has a corresponding reverse complement 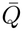, and a node *u* preceding *Q* has a corresponding reverse complement node ū that succeeds 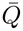. Hence, the number of rows of the interval of 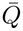 that store each ū, together with the number of rows that store $, determines every subinterval [*s*_*u*_,*e*_*u*_] [18].

The FastLocate structure of the GBWT, an implementation of the r-index locate [32], associates a row with its indexed path *P* and the position *j* of the represented visit to the final node of *Q*. The corresponding occurrence is then (*P, i*), where *i* = *j* − |*Q*| + 1.

### Maximal exact matches

Consider two distinct occurrences (*P, i*) ≠ (*P* ^*′*^, *i*^*′*^) of identical subpaths *P* [*i*..*j*] = *P* ^*′*^[*i*^*′*^..*j*^*′*^], where *P* = (*v*_1_, …, *v*_|*P* |_) and 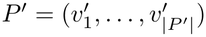 are indexed paths. A match is treated as an unordered pair of occurrences. These occurrences form a *graphical maximal exact match* (gMEM) if they cannot be extended by matching nodes in either direction along their respective paths. Here, exact matching means equality of oriented-node sequences, and match endpoints occur at node boundaries. Thus, the definition does not include matches that begin or end within a node or matches between distinct nodes with identical sequence labels.

Formally, the occurrences constitute a gMEM if both non-extension conditions hold:

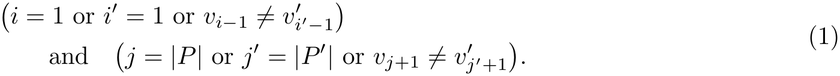

Equivalently, their preceding nodes differ or are both $, and their following nodes differ or are both $. This definition is pairwise, and we call such a match *pairwise-maximal* : *P* [*i*..*j*] need not be maximal in all its occurrences, but only relative to the specific pair of occurrences under consideration.

The general definition permits distinct occurrences on the same indexed path, as well as occurrences on the two orientations of the same input haplotype. For IBD detection between distinct input haplotypes, we retain only matches whose occurrences belong to different input haplotype identifiers.

Because both orientations are indexed, a gMEM between occurrences (*P, i*) and (*P* ^*′*^, *i*^*′*^), ending at positions *j* and *j*^*′*^, has a reverse complement between (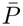, |*P*| − *j* + 1) and (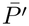, |*P* ^*′*^| − *j*^*′*^ + 1). We regard a match and its reverse complement as equivalent. This equivalence class contains at most two unordered occurrence pairs, allowing for cases in which a match coincides with its reverse complement.

### Maximal subpaths and nodes

Two distinct occurrences of identical subpaths *P* [*i*..*j*] = *P* ^*′*^[*i*^*′*^..*j*^*′*^] are *right-maximal* when their following nodes differ or are both $, formally *j* = |*P*| or *j*^*′*^ = |*P* ^*′*^| or 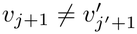 Each pair of corresponding nonempty suffixes of equal node count is then also right-maximal.

We call an oriented node *v* ∈ *V* ^*±*^ a *right-maximal node* if the single-node subpath (*v*) has two distinct occurrences that are right-maximal.

Symmetrically, two distinct occurrences of identical subpaths are *left-maximal* when their preceding nodes differ or are both $, formally *i* = 1 or *i*^*′*^ = 1 or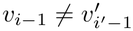. Every pair of corresponding nonempty prefixes of equal node count is then also left-maximal. We call an oriented node *v* ∈ *V* ^*±*^ a *left-maximal node* if the single-node subpath (*v*) has two distinct occurrences that are left-maximal.

Because each path is indexed along with its reverse complement, a predecessor *u* of *v* corresponds to a successor 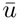 of 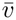, and path starts correspond to path ends under reversal. Hence, *v* is left-maximal if and only if 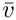 is right-maximal.

## 3 GIBD Algorithm

GIBD identifies segments inherited from a common ancestor, termed identity-by-descent (IBD) segments, across all pairs of haplotypes in a pangenome graph and annotates the structural variants carried by both haplotypes within each segment (Figure 1). To do this, GIBD first traverses the GBWT to identify gMEMs: maximal subpaths traversed identically by two haplotypes (Sections 3.2 and 3.3), and then merges nearby gMEMs between the same haplotype pair into candidate IBD segments. These segments are then annotated with the insertions, deletions, and inversions carried by both haplotypes (Section 3.4).

**Figure 1:**
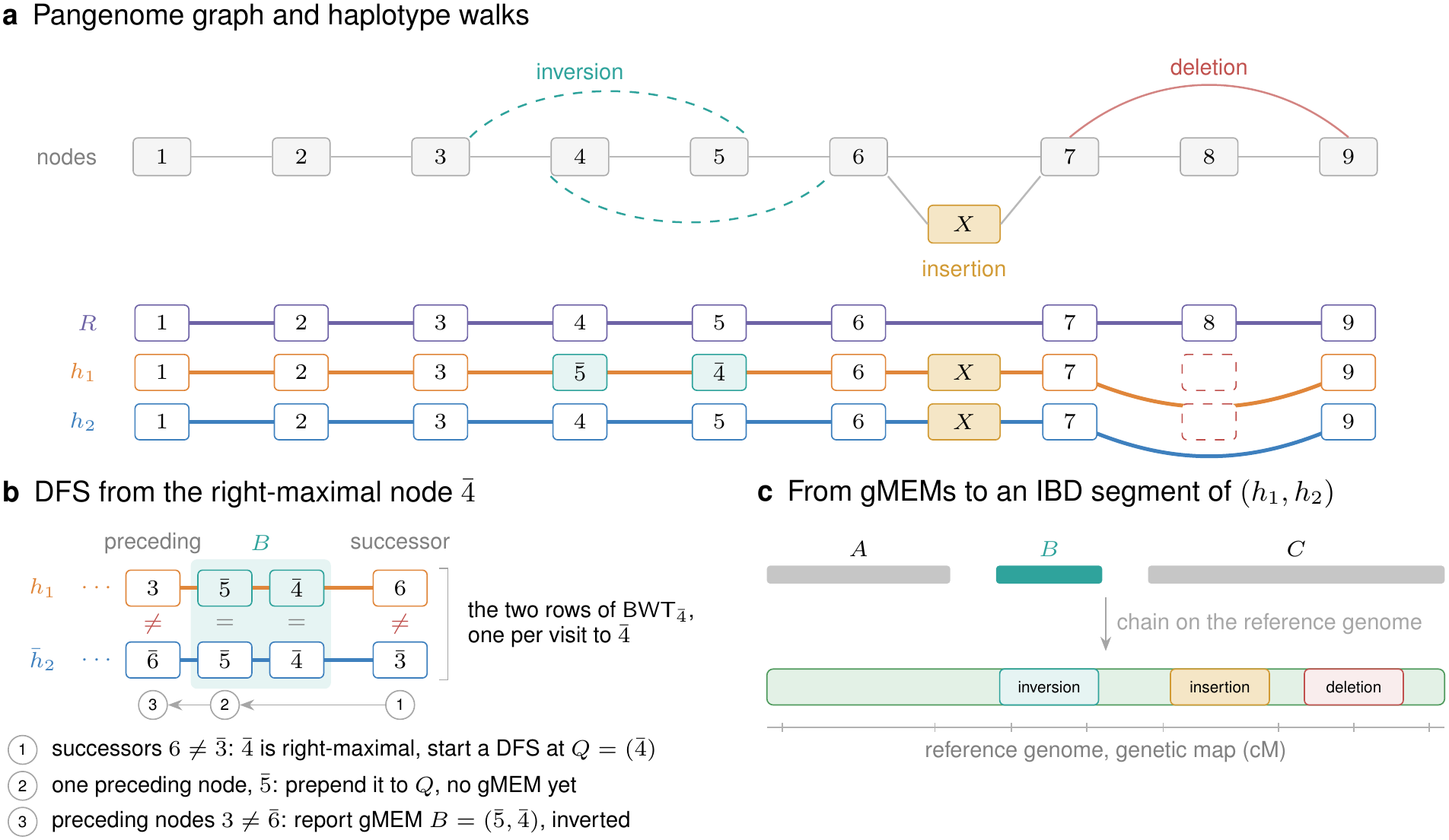
Overview of GIBD. (a) A pangenome graph with the path *R* of the reference genome and two haplotypes. Both carry the insertion *X* and the deletion of node 8, and *h*_1_ carries an inversion of nodes 4 and 5. (b) A depth-first search (DFS) in the GBWT of *h*_1_, *h*_2_ and their reverse complements. (1) BWT 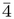 has two rows with different successors, so a DFS starts at 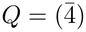. (2) Both occurrences of *Q* are preceded by 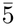, which GIBD prepends. (3) The occurrences of 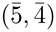 have different preceding nodes and form the inverted gMEM *B*. DFSs from other right-maximal nodes give the gMEMs *A* and *C*. (c) GIBD joins *A, B* and *C* into one IBD segment, labeled as differing by the inversion and with the shared insertion and deletion.

### 3.1 Input to GIBD

GIBD takes as input a GBWT index of the haplotype paths in a pangenome graph, a reference path in the graph, and a genetic map in the coordinates of that reference. GIBD searches the GBWT for gMEMs and retains gMEMs at least *b*_*m*_ base pairs long. It uses the reference coordinates to establish the chromosomal order and spacing of gMEMs. To obtain these coordinates, GIBD walks the reference genome once and records the reference coordinate of each node on it, as the total length of the nodes before it. These coordinates identify where a gMEM shares nodes with the reference and give each gMEM its reference interval (Section 3.4). The corresponding genetic map accounts for variation in recombination rates along the chromosome by converting reference positions of gMEMs to genetic positions in centimorgans (cM).

GIBD merges the gMEMs of each haplotype pair into IBD segments using four length thresholds (Section 3.4). Every IBD segment must contain a gMEM of length at least *s* cM. A gMEM of at least the minimum extension length *x* cM can extend a segment if the gap between the segment and this gMEM is at most *d* base pairs. GIBD reports IBD segments with genetic length at least *c* cM. This threshold helps distinguish recent shared inheritance from chance matches or distant ancestry, since longer genetic segments generally indicate more recent shared ancestry [2]. Finally, GIBD compares both haplotype paths with the reference and annotates shared insertions, deletions, and inversions of at least *b*_*v*_ base pairs, allowing their lengths on the two haplotypes to differ up to a length tolerance threshold of *f* and *f*_*bp*_.

### 3.2 Left-extension of right-maximal nodes

To enumerate gMEMs, GIBD first identifies right-maximal nodes by traversing the GBWT and stores them on a stack. Next, GIBD pops a node *v* from the stack and initiates a depth-first search from the single-node subpath *Q* = (*v*). The occurrences of *Q* correspond to the rows of BWT_*v*_. The search extends *Q* to the left one node at a time. These extensions preserve *v* as the rightmost node, so their occurrences correspond to intervals [*s, e*] in BWT_*v*_. For each possible left-extension node *u*, the occurrences of *Q* that can be extended to *uQ* correspond to a subinterval [*s*_*u*_, *e*_*u*_] ⊆ [*s, e*] in BWT_*v*_.

To extend *Q* to the left, GIBD retrieves these subintervals from the interval of the reverse complement subpath 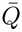. It does this with a successor lookup in 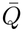, as the successors of subpath 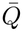 are the reverse complements of the predecessors of subpath *Q*. GIBD traverses the extension *uQ* only if two of its occurrences form a right-maximal pair. This holds if two rows of the subinterval store different successors, or if two rows both store $, so that both occurrences end at a path end. Otherwise, no extension of *uQ* is a gMEM, and GIBD prunes the branch. If all occurrences of *Q* share the same preceding node *u*, then no pair is left-maximal, so *u* is prepended to *Q* and the extension of *Q* continues. If occurrences of *Q* have distinct preceding nodes, then at least one pair is left-maximal. In this case, GIBD enumerates all pairs that form gMEMs (see Section 3.3) and continues along each extension.

### 3.3 Identification of gMEMs

When a left-maximal subpath *Q* is identified from the traversal, GIBD identifies the pairs of occurrences that form gMEMs. By Equation (1), two occurrences of the subpath *Q* with predecessor nodes *u* and *u*^*′*^, and successor nodes *w* and *w*^*′*^ form a gMEM when *u* ≠ *u*^*′*^ or *u* = *u*^*′*^ = $, and *w* ≠ *w*^*′*^ or *w* = *w*^*′*^ = $. Whether a pair forms a gMEM therefore depends only on its four boundary nodes. Thus, GIBD evaluates this condition once for each distinct combination of preceding and following nodes (*u, w, u*^*′*^, *w*^*′*^) between pairs of occurrences. For each combination satisfying the condition, GIBD uses FastLocate to recover each occurrence as a pair (*P, i*), where *P* is the indexed haplotype path containing *Q* and *i* is its starting position within *P* . Then it enumerates each gMEM as a pair of occurrences (*P, i*), (*P* ^*′*^, *i*^*′*^) of *Q*, where the first occurrence is flanked by (*u, w*) and the second by (*u*^*′*^, *w*^*′*^). Each enumerated pair is left- and right-maximal. The traversal enumerates each gMEM exactly once before filtering; see Theorem 1. Since the GBWT indexes both orientations of each haplotype path, a gMEM and its reverse complement are enumerated as distinct gMEMs. GIBD discards one gMEM of each such pair (Appendix A). It also discards gMEMs shorter than *b*_*m*_ bp, and gMEMs whose occurrences lie on the same haplotype. When repeated occurrences of *Q* produce multiple gMEMs between the same pair of haplotypes in the same relative orientation, GIBD retains only one of them.

### 3.4 From gMEMs to IBD segments

Mutations and sequencing errors can break exact matches within an IBD segment, leaving multiple shorter gMEMs. GIBD orders the gMEMs along the chromosome and merges them into IBD segments [12]. To order gMEMs along the chromosome and estimate the genomic distance between them, GIBD identifies the nodes each gMEM shares with the reference path. This can be done in *O*(1) time per node using the auxiliary data structure that maps each node to the reference. If the gMEM does not share any nodes with the reference, then GIBD discards it. If it shares two or more nodes, GIBD retrieves their coordinates from the auxiliary data structure and assigns the gMEM the interval from the smallest coordinate to the end of the node with the largest coordinate. When a gMEM shares exactly one node with the reference, GIBD traces the alternate allele along the haplotype path, starting at each end of the gMEM that is not on the reference. If the path rejoins the reference, GIBD assigns the gMEM the interval between the shared node and the point of reentry. Otherwise, GIBD assigns the gMEM the interval of the shared node. The genetic map converts the two ends of each interval to cM, and their difference is the genetic length of the gMEM. After placing the gMEMs on the reference, GIBD discards gMEMs whose assigned intervals span less than *x* cM. GIBD handles gMEMs that the two haplotypes traverse in opposite orientations separately (see *Inversion on one haplotype* below). GIBD sorts the remaining gMEMs by the left endpoint of their reference intervals and scans them in that order. The first gMEM initializes a candidate segment. For each subsequent gMEM, GIBD adds it to the current candidate if its interval starts at most *d* bp beyond the candidate’s right endpoint. After each addition, the candidate interval spans all gMEMs assigned to it. Otherwise, the candidate closes and the next gMEM initializes a new one.

GIBD reports a candidate as an IBD segment only if it spans at least *c* cM and contains at least one gMEM spanning *s* cM or more. If the two haplotypes belong to the same individual, GIBD reports the segment as homozygosity by descent (HBD).

Next, GIBD identifies SVs on each haplotype within an IBD segment. For each haplotype, GIBD orients its path relative to the reference and uses the reference nodes that the haplotype traverses in this orientation as anchors. Between consecutive anchors, GIBD calls an insertion or a deletion if the haplotype sequence is at least *b*_*v*_ bp longer or shorter than the corresponding reference sequence, respectively. It calls an inversion if the haplotype traverses reference nodes in the opposite orientation between the two anchors and those nodes have a combined length of at least *b*_*v*_ bp. GIBD records the type, reference interval, and length of each call. Two calls, one from each haplotype, are considered the same SV if they have the same type, their reference intervals overlap, and their lengths *L*_1_ and *L*_2_ (in bp) satisfy |*L*_1_ −*L*_2_| ≤ max {*f*_*bp*_, *f* max(*L*_1_,*L*_2_)} . GIBD annotates the IBD segment with the resulting shared SVs. The length tolerance accommodates small differences in the inferred length of the same variant due to assembly errors [33].

#### Inversion on one haplotype

Local orientation differences can also occur between paired haplotypes. For example, if one haplotype traverses the node path (*u, v, w*) and the other traverses (*u*, 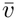, *w*), the haplotypes share a gMEM ending at *u*, an inverted gMEM spanning *v*, and a gMEM beginning at *w*. Because the haplotypes differ only in the local orientation of this sequence, GIBD merges the inverted gMEM with its two flanking gMEMs into a single segment, even if the inverted gMEM is shorter than *x* cM, provided that the intervening gaps do not exceed *d* base pairs.

## 4 Correctness and Complexity Guarantees

We first establish the correctness of the GIBD traversal and then state its running time.

### Theorem 1.

*Let the GBWT index the haplotype paths of the input graph, and let* (*u, w, u*^*′*^, *w*^*′*^) *be the boundary nodes of a pair of occurrences of a subpath Q in those paths. The GIBD traversal enumerates the pair if and only if it forms a gMEM, and enumerates each such pair exactly once*.

#### Proof.

For every pair of occurrences enumerated by GIBD, the corresponding boundary nodes (*u, w, u*^*′*^, *w*^*′*^) satisfy Equation (1). By definition, each enumerated pair therefore forms a gMEM. Conversely, let *p* and *p*^*′*^ be two occurrences of *Q* that form a gMEM, and let *v* be the rightmost node of *Q*. By right maximality, *p* and *p*^*′*^ are followed by distinct nodes or both end at path boundaries. The same condition holds for the corresponding occurrences of every suffix of *Q*, including (*v*), because each suffix ends at the same position as *Q* in each occurrence. Thus, GIBD starts a DFS at *v* and follows the sequence of left extensions from (*v*) to *Q*, since each intermediate suffix has a right-maximal pair. Once the DFS reaches *Q*, GIBD checks the boundary nodes of *p* and *p*^*′*^. The pair satisfies Equation (1) by assumption, so GIBD enumerates it.

For uniqueness, let *Q* = (*v*_1_, …, *v*_*k*_). Since each DFS step prepends one node, the only way to reach *Q* is to start at (*v*_*k*_) and prepend *v*_*k−*1_, …, *v*_1_ in that order. At *Q*, each occurrence has a unique ordered pair of boundary nodes. Thus, an unordered pair of distinct occurrences determines a unique combination of two such pairs of boundary nodes, which may be identical. GIBD examines each combination once and enumerates each qualifying pair of occurrences within it once. Hence, each gMEM is enumerated exactly once. □

Let *n* be the total number of node occurrences across all haplotype paths indexed by the GBWT. Each visit to a node contributes one to this count, and both orientations of every haplotype path are included. Let *occ* be the number of gMEMs enumerated before filtering, counting each qualifying pair of subpath occurrences separately.

On a graph of bounded degree, GIBD uses a balanced move structure to perform each successor lookup in *O*(1) time [34, 35]. For each DFS root *v*, GIBD locates each row of BWT_*v*_ once, in *O*(1) time per row [32, 36].

### Theorem 2.

*If G has bounded degree, the GIBD traversal visits each subpath at most once and enumerates all occ gMEMs in O*(*n* + *occ*) *time*.

This bound is linear in the number of node visits on the indexed paths and the number of gMEMs enumerated. The proof of Theorem 2 is in Appendix A, together with the space bounds of GIBD.

## 5 Results

### 5.1 Experimental setup

#### Implementation

GIBD is implemented in C++17 and available at https://github.com/uwaiseibna/GIBD, with GBWT and sdsl-lite as dependencies [18, 37]. GIBD implements balanced move structures over the GBWT for its bidirectional and locate queries [34, 35]. It was compiled with GCC 11.5 at `-O2`. Unless stated otherwise, all methods ran single-threaded on an AMD EPYC 7702 node with 1 TB of memory, and we measured wall-clock time and peak memory with GNU `time`. The time and memory of GIBD include SV annotation. We report memory in GB of 2^30^ bytes.

#### Competing methods

We compare GIBD with the pangenome-based IBD method `impop`_*k*_ [20]. Because there are few pangenome-based IBD methods, we also compare GIBD with the linear IBD methods `hap-ibd` [12], `RaPID` [13], `iLASH` [14] and `TPBWT` [15]. We also compare the gMEMs of GIBD with the matches reported by `GBWT-Query` [19].

#### Metrics and parameters

We evaluate IBD detection using four metrics. *Power* is the average fraction of each known IBD segment covered by its best-overlapping reported segment between the same haplotypes. *Accuracy* is the fraction of reported segments that overlap a known IBD segment between the same haplotypes. *Detected* is the fraction of known IBD segments overlapped by a reported segment between the same haplotypes. *Length accuracy* is the fraction of the total length of reported segments that overlaps known IBD segments between the same haplotypes. For SV annotation, *precision* is the fraction of reported SV labels that are correct, and *recall* is the fraction of known SV labels recovered.

GIBD uses *s* = 2 cM, *x* = 1 cM, *c* = 2 cM, *b*_*m*_ = 1,000 bp, *b*_*v*_ = 50 bp, *f*_*bp*_ = 10 bp, *f* = 0.1, and *d* = 1,000 bp. For SVs of at least *b*_*v*_ base pairs, the length tolerance is at most 20% of the SV length, stricter than the 30% tolerance of Truvari [33]. Each experiment uses the reference genome that its truth or competing method requires, namely GRCh37 for the benchmark of linear methods, T2T-CHM13 for the comparison with `impop`_*k*_ and scaling, and GRCh38 for SV annotation.

### 5.2 Validation and comparison of gMEMs

We first verified that GIBD reports exactly the gMEMs of a GBWT. `GBWT-Query` [19] reports long matches between one query haplotype at a time and the haplotypes indexed in a GBWT. Since `GBWT-Query` cannot extend these matches or identify IBD segments, our comparison is limited to gMEM detection.

Following `GBWT-Query`’s evaluation, we compared it to GIBD on the GBWT of 1000 Genomes Project chromosome 21 containing 5,008 haplotypes. `GBWT-Query` builds its index on 4,908 of these haplotypes and uses the remaining 100 as queries. For each query haplotype, it reports the long matches between the query and the indexed haplotypes. GIBD instead ran once on the full GBWT and found the gMEMs between all pairs of haplotypes. For the comparison, we kept only the gMEMs between a query haplotype and an indexed haplotype. We evaluated matches at two minimum match length thresholds of 7,500 and 20,000 nodes. GIBD and `GBWT-Query` reported identical sets of 631,382 matches at 7,500 nodes and 491,724 matches at 20,000 nodes, with identical haplotypes, start positions and lengths across all 100 queries.

### 5.3 Comparison to linear IBD methods

We next compared GIBD with linear methods on simulated data with known IBD. Linear methods cannot annotate structural variants, so this experiment evaluates only how well GIBD detects IBD segments compared with linear methods.

We used the chromosome 20 dataset simulated by Tang et al. [26], comprising 1,000 individuals generated under an out-of-Africa demographic model. We used GRCh37 and its genetic map to maintain consistency with the simulation. The linear methods read the VCF of the simulation, and GIBD read a graph with one bubble per site of the same VCF. We evaluated the methods in eight experiments, combining two site sets, all sites or sites with minor allele frequency (MAF) ≥ 0.1, with four genotyping error rates of 0, 0.0125%, 0.05% and 0.1%. `IBD_BM` [26] evaluated each method against the known segments, stratified by known segment length (Appendix B).

GIBD achieved power and accuracy comparable to those of `hap-ibd`, with absolute differences of at most 0.009 and 0.001, respectively, across all eight experiments and segment-length bins (Figure 2a). For 2–3 cM segments, both methods maintained accuracy of at least 0.994. `RaPID` and `TPBWT` achieved near-complete power on these segments, but their accuracy fell as low as 0.079 and 0.275, respectively. `iLASH` maintained similarly high accuracy to GIBD and `hap-ibd`, but generally had lower power. Because GIBD and `hap-ibd` construct IBD segments from exact matches, their power decreased with increasing genotyping error, whereas `RaPID` and `TPBWT` largely retained their power.

**Figure 2:**
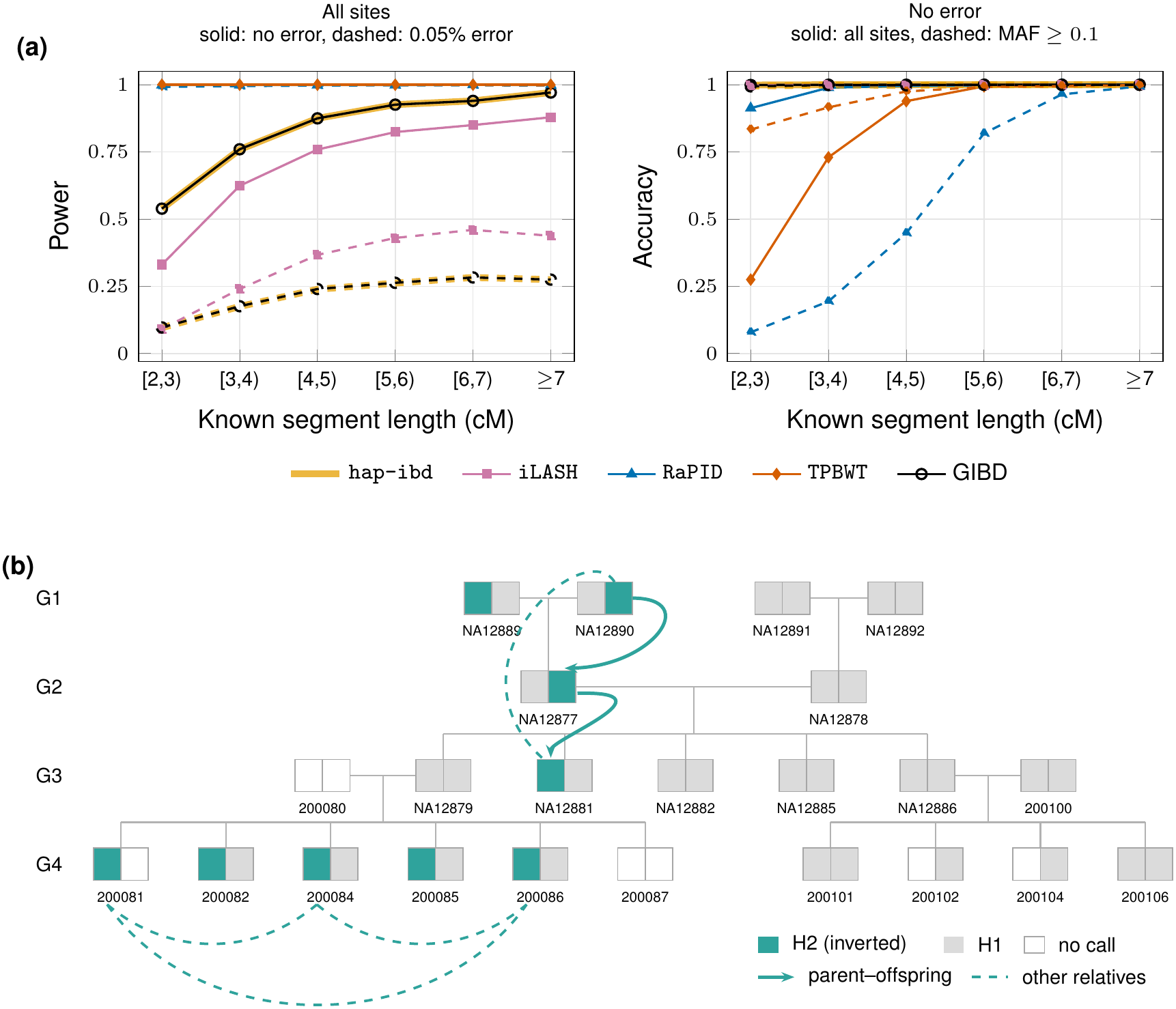
(a) Power and accuracy of GIBD and the linear methods on the benchmark of Tang et al. [26] by known segment length (all eight experiments in Appendix B). (b) The 17q21.31 inversion in the Platinum Pedigree, generations G1–G4. Each box is one member, with its two haplotypes (paternal left and maternal right from G2 on) colored by H1 or H2. Arrows (parent–offspring) and dashed arcs (other relatives) mark the segments that GIBD labeled with the H2 inversion at seed and extension lengths of 1 and 0.5 cM.

Table 1 summarizes the trade-off between power and accuracy on 2–3 cM segments by their harmonic mean, the F1 score. No method achieved the highest F1 score in all eight experiments. `RaPID` had the highest F1 score in the four all-sites experiments. Among the experiments restricted to sites with MAF ≥ 0.1, `hap-ibd` had the highest F1 score without genotyping error, GIBD at an error rate of 0.0125%, and `TPBWT` at error rates of 0.05% and 0.1%. GIBD was slower and used more memory than the linear methods (Appendix B).

**Table 1:** F1 score (harmonic mean of power and accuracy) on known segments of 2–3 cM in the eight experiments on the simulation of Tang et al. [26]. The highest value of each experiment is in bold.

| Sites | Error | GIBD | hap-ibd | iLASH | RaPID | TPBWT |
|---|---|---|---|---|---|---|
| All | 0 | 0.700 | 0.700 | 0.497 | **0.954** | 0.431 |
|  | 0.0125% | 0.494 | 0.492 | 0.373 | **0.954** | 0.438 |
|  | 0.05% | 0.177 | 0.175 | 0.167 | **0.951** | 0.455 |
|  | 0.1% | 0.051 | 0.049 | 0.062 | **0.944** | 0.480 |
| MAF ≥ 0.1 | 0 | 0.996 | **0.997** | 0.911 | 0.146 | 0.903 |
|  | 0.0125% | **0.953** | 0.952 | 0.866 | 0.146 | 0.904 |
|  | 0.05% | 0.839 | 0.836 | 0.747 | 0.146 | **0.907** |
|  | 0.1% | 0.710 | 0.703 | 0.615 | 0.146 | **0.909** |

### 5.4 Comparison to a pangenome-based IBD method

Next, we compared GIBD to `impop`_*k*_, a pangenome-based IBD method. Since `impop`_*k*_ does not annotate structural variants, we compared only IBD detection.

We simulated haplotypes with known IBD from HPRC v2 assemblies [27] on T2T-CHM13 [38], the reference genome of `impop`_*k*_. As founders, we used haplotype walks from the HPRC v2 Minigraph-Cactus graph spanning chr22:21.7–38 Mb, the longest interval on chromosome 22 for which at least 400 HPRC v2 haplotypes are assembled as a single contiguous walk. We excluded walks containing a somatic V–J deletion at the IGL locus [39]. We simulated ancestry using `msprime` [40], with a single population growing exponentially from 100 diploid individuals 40 generations ago to 2,000 at present. We sampled 500 diploid individuals and modeled recombination using the T2T-CHM13 genetic map [41]. Each ancestral lineage present 40 generations ago was assigned a distinct founder walk.

`impop`_*k*_ compares every pair of haplotypes in every window, so we first measured how it scales against GIBD as the length of the graph grows (Table 2). We used nested regions of 5, 10 and 15 Mb and the whole simulated region of 16.3 Mb from 200 haplotypes of 100 randomly chosen individuals. The time of GIBD includes building the GBWT and IBD detection, and the time of `impop`_*k*_ excludes the alignment of each haplotype to the reference prior to detection [42]. GIBD took 33 s and 0.9 GB on 5 Mb and 133 s and 2.1 GB on the whole region. `impop`_*k*_ took 9.6 h and 6.8 GB on 5 Mb and 38.2 h and more than 64 GB on the whole region. On the 15 and 16.3 Mb regions, one 10 kb window (chr22:24.08–24.09 Mb) needed more than the 64 GB memory limit, so its `impg` process was stopped and `impop`_*k*_ skipped that window. Next, we compared the accuracy in IBD detection on the 5 Mb region chr22:30–35 Mb (5.7 cM).

**Table 2:**
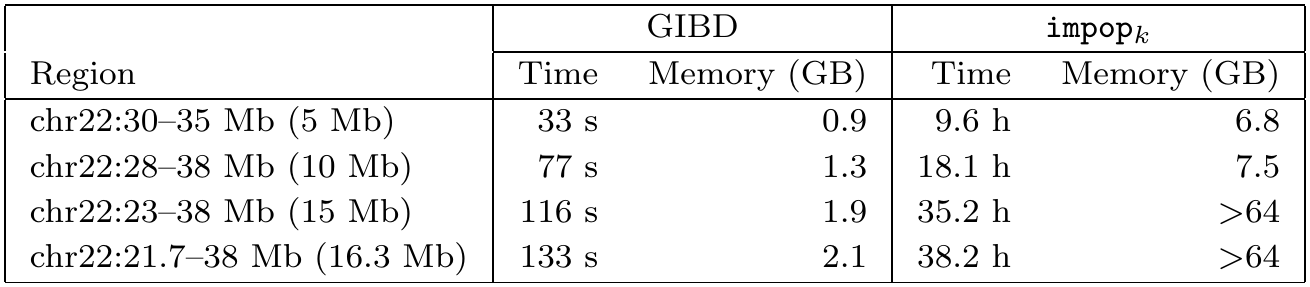
Wall-clock time and peak memory of GIBD and impop_*k*_ on the same 200 simulated haplotypes, by length of the T2T-CHM13 chr22 region.

|  | GIBD |  | impop<sub>*k*</sub> |  |
|---|---|---|---|---|
| Region | Time | Memory (GB) | Time | Memory (GB) |
| chr22:30–35 Mb (5 Mb) | 33 s | 0.9 | 9.6 h | 6.8 |
| chr22:28–38 Mb (10 Mb) | 77 s | 1.3 | 18.1 h | 7.5 |
| chr22:23–38 Mb (15 Mb) | 116 s | 1.9 | 35.2 h | >64 |
| chr22:21.7–38 Mb (16.3 Mb) | 133 s | 2.1 | 38.2 h | >64 |

`IBD BM` evaluated both methods against the known segments between the 100 individuals. Both methods had accuracy of at least 0.98 in every length bin (Table 3). GIBD detected more short segments, 0.61 and 0.90 in the 2–3 and 3–4 cM bins against 0.52 and 0.79, partly because the 2 Mb threshold of `impop`_*k*_ corresponds to 2.3 cM in this region. GIBD had higher length accuracy, 0.97–0.99 against 0.85–0.97. Both methods detected every segment of 4–6 cM. From 3 cM on, `impop`_*k*_ covered more of each segment (power 0.79–1.00 against 0.77–0.95), since its identity model tolerates mismatches, whereas a segment of GIBD ends at its last exact match.

**Table 3:** Accuracy, detection rate, power and length accuracy of GIBD and impop_*k*_ on 200 simulated haplotypes in T2T-CHM13 chr22:30–35 Mb, by known segment length (cM).

|  |  | known segment length (cM) |  |  |  |
|---|---|---|---|---|---|
| Metric | Method | [2, 3) | [3, 4) | [4, 5) | [5, 6) |
| Number of known segments |  | 399 | 303 | 38 | 166 |
| Accuracy | GIBD | 0.995 | 1.000 | 1.000 | 1.000 |
|  | impop<sub>*k*</sub> | 0.992 | 0.991 | 1.000 | 0.984 |
| Detected | GIBD | 0.614 | 0.898 | 1.000 | 1.000 |
|  | impop<sub>*k*</sub> | 0.516 | 0.792 | 1.000 | 1.000 |
| Power | GIBD | 0.591 | 0.773 | 0.946 | 0.918 |
|  | impop<sub>*k*</sub> | 0.515 | 0.790 | 0.998 | 1.000 |
| Length accuracy | GIBD | 0.987 | 0.985 | 0.971 | 0.977 |
|  | impop<sub>*k*</sub> | 0.850 | 0.974 | 0.902 | 0.950 |

### 5.5 Annotating IBD segments with structural variants

We then tested whether GIBD annotates each IBD segment with the correct SVs. We used GRCh38 as the reference genome for all SV annotation, since the inheritance truthset and the SV callset of the Platinum Pedigree, a real multi-generation family, are defined on GRCh38. We first evaluated the SV labels on simulated and real haplotypes, and then traced a disease-associated SV on real haplotypes across generations with GIBD.

#### Simulated haplotypes

We adopted the simulation approach of Section 5.4 on the HPRC v2 graph built on GRCh38 and generated two simulated graphs, one without and one with assembly error (Appendix C). Of the IBD segments GIBD detected, its insertion and deletion labels had precision and recall of 1.000, and its inversion labels had precision 1.000. When exactly one of the two haplotypes carries an inversion, GIBD labeled every segment that spans it as differing by the inversion, and labeled none of the about 119,700 pairs that do not differ. Assembly error lowered detection but not precision (Table A3).

#### Four-generation pedigree

We then evaluated SV annotation of GIBD in a real family with known inheritance. We used the phased assemblies of the 23 openly released members of the Platinum Pedigree, the four-generation family CEPH 1463 (46 haplotypes) [28, 43], and built a pangenome graph of chromosome 20, consistent with the linear benchmark (Section 5.3), with GRCh38 as the reference genome [44]. We derived the known inheritance of each child haplotype from the consortium’s phased family truthset, independently of the assemblies. Each child inherits one of the two haplotypes of each parent, so the SVs at which a parent is heterozygous show whether an IBD segment links the child to the correct parental haplotype. We compared GIBD’s SV labels with the consortium’s SV callset.

GIBD’s IBD detection had perfect accuracy. At seed and extension lengths of 1 and 0.5 cM, parent– offspring segments mapped each child to the correct parental haplotype at all 3,235 heterozygous-parent SVs inside its segments (Appendix D). Power was 0.61–0.87, because assembly differences between inherited haplotypes still break some exact matches. Against the callset, GIBD’s insertion labels had precision 0.57–0.58 and recall 0.62–0.66, and its deletion labels had precision 0.70–0.73 and recall 0.68–0.78. GIBD reads SVs from the pangenome graph, and the graph does not carry every SV in the callset. On single haplotypes, the graph’s SVs had precision 0.53 and recall 0.62 for insertions and precision 0.63 and recall 0.63 for deletions. Most SVs the graph misses lie in tandem repeats (88% of insertions and 81% of deletions), where the graph often gives the repeat a different length than the callset. When the graph carries a callset SV on both haplotypes of a covered known segment, GIBD labeled it for 99.7% of insertions and all deletions.

To trace a disease-associated inversion, a pangenome graph of chromosome 17 was built from the same assemblies. The 900-kb inversion at 17q21.31 separates the H1 and H2 haplotypes [7], and H2 carries the inverted orientation relative to GRCh38. This region is a known disease risk locus [29, 30]. Since the callset contains only SVs up to 100 kb, we determined whether each haplotype carries H1 or H2 directly from its assembly alignment to GRCh38 [42]. NA12877 inherited H2 from NA12890 and passed it to NA12881, and five children of 200080 carry it on their paternal haplotype (Figure 2b). At seed and extension lengths of 1 and 0.5 cM, 23 GIBD segments between relatives span the inversion. GIBD labeled all 6 segments between two H2 haplotypes with the shared inversion, including the two parent–offspring transmissions (arrows in Figure 2b), and all 17 segments between two H1 haplotypes with the reference orientation.

### 5.6 Scalability in time and memory

Finally, we measured how the time and memory of GIBD grow with the number of haplotypes on whole chromosomes. We ran GIBD on chromosomes 1 and 22 of the HPRC v2 graph built on T2T-CHM13 [27] with the corresponding map [41]. Chromosome 22 spans 51.3 Mb and 4.1 million nodes, and chromosome 1 spans 248.4 Mb and 10.9 million nodes.

On chromosome 22, GIBD built the GBWT and detected IBD for 58, 116, 232 and 462 haplotypes in 6.5, 13.6, 29.1 and 50.0 min, with at most 3.9, 5.5, 8.6 and 14.9 GB of memory (Table 5). On chromosome 1, it took 159.2 min and at most 54.1 GB for 462 haplotypes. From 58 to 462 haplotypes, an 8 × increase, time grew 7.7× on chromosome 22 and 11.9× on chromosome 1. Memory increased 3.8 and 4.4 times. In the same range, the number of gMEMs increased about 50 times, from 0.12M to 5.70M on chromosome 22 and from 0.46M to 22.98M on chromosome 1.

**Table 4:**
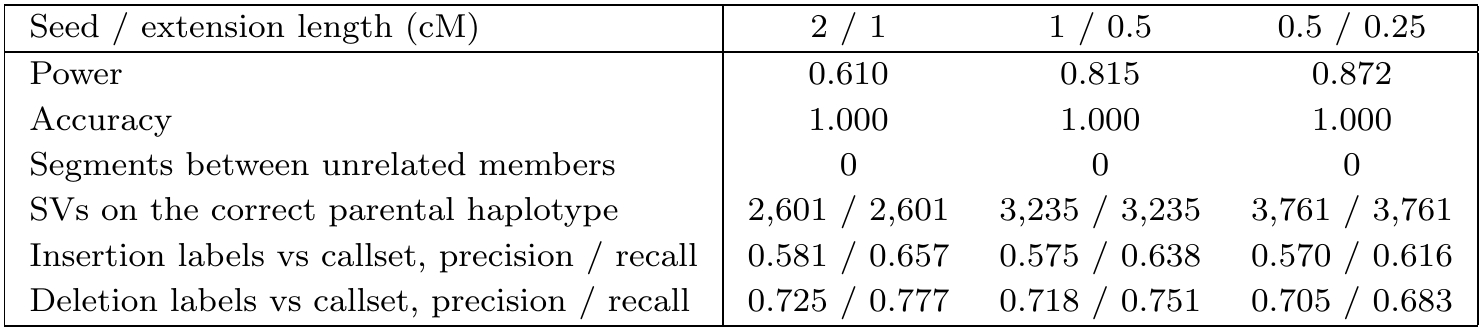
GIBD in the Platinum Pedigree, chromosome 20, on the 54 known segments of at least 2 cM. Labels are compared with the callset using Truvari [33].

**Table 5:** Wall-clock time (min) and peak memory (GB) of building the GBWT and of the search, on chromosomes 1 and 22 of the T2T-CHM13 HPRC v2 graph. gMEMs, in millions.

|  | Chromosome 22 |  |  |  |  | Chromosome 1 |  |  |  |  |
|---|---|---|---|---|---|---|---|---|---|---|
|  | Build |  | Search |  |  | Build |  | Search |  |  |
| Haplotypes | Min | GB | Min | GB | gMEMs | Min | GB | Min | GB | gMEMs |
| 58 | 2.6 | 3.4 | 3.9 | 3.9 | 0.12 | 5.3 | 10.6 | 8.0 | 12.3 | 0.46 |
| 116 | 5.5 | 4.6 | 8.0 | 5.5 | 0.33 | 11.0 | 15.3 | 17.2 | 18.4 | 1.81 |
| 232 | 11.7 | 7.1 | 17.4 | 8.6 | 1.46 | 26.4 | 24.5 | 40.9 | 30.4 | 6.34 |
| 462 | 19.3 | 11.9 | 30.7 | 14.9 | 5.70 | 59.4 | 42.9 | 99.8 | 54.1 | 22.98 |

## 6 Conclusion

GIBD is, to our knowledge, the first method that detects IBD in a pangenome graph and labels each segment with its insertions, deletions and inversions, even an inversion that only one haplotype carries. It matched `hap-ibd` in power and accuracy and found more short segments than `impop`_*k*_ in 33 seconds instead of 9.6 hours. Its SV labels had precision 1.000 in simulation, and in a real family its segments followed 3,235 SVs and the 17q21.31 inversion across generations. IBD now reports which structural variants were inherited, not only where.

