## supplement for "Structural Variant Aware Identity-by-Descent Detection on Pangenome Graphs"

#### A Reverse complements, time and space of the traversal

The GBWT indexes every haplotype path in both orientations. GIBD needs the reverse orientation to find inverted gMEMs, which mark inversions. As a result, the reverse complement of every gMEM is also a gMEM, and the traversal finds both.

**Lemma A1.** *The traversal finds each gMEM together with its reverse complement, and GIBD keeps exactly one of the two.*

*Proof.* Let a subpath $Q = (v_1, \ldots, v_k)$ occur on paths $P$ and $P'$ as a gMEM. Read backwards, $Q$ is its reverse complement $\bar{Q} = (\bar{v}_k, \ldots, \bar{v}_1)$, which occurs on $\bar{P}$ and $\bar{P}'$. The traversal finds the gMEM in the DFS from $v_k$, the last node of $Q$, and its reverse complement in the DFS from $\bar{v}_1$, the last node of $\bar{Q}$.

The GBWT numbers the input haplotypes. Of the two haplotypes of $P$ and $P'$, let $h$ be the one with the smaller number. The reverse complement reads both paths in the opposite orientation of the gMEM. Hence, $h$ is read forward in exactly one of the two. GIBD keeps that one and discards the other. Both cover the same nodes of the same two haplotypes, so GIBD loses no gMEM. □

*Proof of Theorem 2.* The bound covers the traversal and excludes building the GBWT, FastLocate and the move structures. GIBD first reads each GBWT record once to find the right-maximal nodes. Each row of a record is one node visit on one path, so the records have $n$ rows in total. Every node of the GBWT lies on an indexed path, and the header of its record lists only the outgoing edges that the paths use. Hence, the headers have total size $O(n)$, and finding the roots takes $O(n)$ time.

A DFS from a root $v$ visits only subpaths that end at $v$, so two DFSs never visit the same subpath. Inside one DFS, a subpath is reached only from the subpath that is one node shorter on the left. Hence, the DFS reaches each subpath along one branch, and visits each subpath at most once.

The DFS visits a subpath only if two of its occurrences have different succeeding nodes or both end at a path boundary. Give each indexed path its own end symbol. Then every visited subpath has two occurrences followed by different symbols. Hence, every visited subpath is a branching node of the generalized suffix tree of the indexed paths. This tree has $O(n)$ branching nodes, so the DFS visits $O(n)$ subpaths.

Before the DFS from $v$, GIBD indexes the runs of $\mathsf{BWT}_v$. For each row, it stores the run that holds the row. For each run and each succeeding node, it stores the next run with that succeeding node. Since $G$ has bounded degree, this takes time linear in the length of $\mathsf{BWT}_v$, and $O(n)$ time over all roots. With this index, GIBD tests in $O(1)$ time whether a range of rows has two different succeeding nodes, and finds which succeeding nodes occur in the range. At every $B$-th run, for a constant $B$, the move structure stores the nearest runs with each successor. These range samples give the distinct successors of a range of rows in $O(1)$ time. For the range of $\bar{Q}$, these successors are the reverse complements of the distinct preceding nodes of $Q$. Each left extension is then one successor lookup, which takes $O(1)$ time. A visited subpath has $O(1)$ combinations of boundary nodes $(u, w, u', w')$, and GIBD evaluates Equation (1) once for each. Hence, each visit takes $O(1)$ time, and all visits take $O(n)$ time.

In the DFS from $v$, every occurrence of a visited subpath is a row of $\mathsf{BWT}_v$. GIBD locates each row of $\mathsf{BWT}_v$ once, in $O(1)$ time, and reuses the result. The records have $n$ rows in total, so locating takes $O(n)$ time.

For each combination of boundary nodes that satisfies Equation (1), GIBD generates every pair of occurrences, in $O(1)$ time per pair. By Theorem 1, each generated pair is a gMEM and is generated once. Hence, generating the pairs takes $O(occ)$ time. In total, the traversal takes $O(n + occ)$ time. □

The number $occ$ counts every pair that the DFS finds. It includes each gMEM and its reverse complement (Lemma A1), and the pairs that the filters of Section 3.3 later remove. A single subpath can yield a number of pairs quadratic in the number of haplotypes, when two groups $(u, w)$ and $(u', w')$ that satisfy Equation (1) each contain many occurrences.

GIBD processes the DFS roots one at a time. Besides the GBWT, FastLocate and the move structures, it stores three things. The DFS stack grows with the length of the current subpath. The located rows of the current root take at most the size of the largest GBWT record. The pairs generated at the current subpath take space proportional to their number. The GBWT, FastLocate and the move structures take $O(r)$ space for $r$ runs in the GBWT records. These bounds exclude building the index and reporting IBD segments.

### B Comparison to linear IBD methods

Table A1 gives the power and accuracy of every method in the eight experiments of Section 5.3. The linear methods ran with the parameters of the benchmark. Our runs reproduced the calls distributed with the benchmark exactly for `hap-ibd` and `RaPID`, for 99.99% of calls for `TPBWT`, and within the run-to-run variation of `iLASH`. GIBD masks nodes traversed by fewer than two haplotypes, so it used only sites with minor allele count of at least 2, as `hap-ibd` does by default. With singleton sites in the graph, GIBD's power on 2–3 cM segments in the all-sites experiment without error fell from 0.539 to 0.323, because a singleton inside a known segment breaks an exact match.

Table A1: Power (P) and accuracy (A) from `IBD_BM` by known segment length (cM) in all eight experiments on the simulation of Tang et al. [1].

| | | All sites | | | MAF $\geq 0.1$ | | |
|---|---|---|---|---|---|---|---|
| Error | Method | P [2,3) | P $\geq 7$ | A [2,3) | P [2,3) | P $\geq 7$ | A [2,3) |
| 0 | GIBD | 0.539 | 0.971 | 0.999 | 0.999 | 0.999 | 0.994 |
| | `hap-ibd` | 0.539 | 0.971 | 0.999 | 0.999 | 0.999 | 0.995 |
| | `iLASH` | 0.331 | 0.879 | 0.999 | 0.839 | 0.994 | 0.997 |
| | `RaPID` | 0.999 | 0.999 | 0.913 | 0.999 | 0.999 | 0.079 |
| | `TPBWT` | 1.000 | 1.000 | 0.275 | 0.986 | 1.000 | 0.833 |
| 0.0125% | GIBD | 0.328 | 0.733 | 0.999 | 0.915 | 0.982 | 0.994 |
| | `hap-ibd` | 0.326 | 0.733 | 0.999 | 0.913 | 0.982 | 0.995 |
| | `iLASH` | 0.229 | 0.718 | 0.998 | 0.766 | 0.885 | 0.997 |
| | `RaPID` | 0.998 | 0.999 | 0.913 | 0.999 | 0.999 | 0.079 |
| | `TPBWT` | 1.000 | 1.000 | 0.280 | 0.986 | 1.000 | 0.835 |
| 0.05% | GIBD | 0.097 | 0.275 | 1.000 | 0.726 | 0.866 | 0.994 |
| | `hap-ibd` | 0.096 | 0.275 | 1.000 | 0.721 | 0.865 | 0.995 |
| | `iLASH` | 0.091 | 0.438 | 0.998 | 0.597 | 0.682 | 0.997 |
| | `RaPID` | 0.992 | 0.997 | 0.913 | 0.998 | 0.999 | 0.079 |
| | `TPBWT` | 0.999 | 0.998 | 0.295 | 0.986 | 1.000 | 0.839 |
| 0.1% | GIBD | 0.026 | 0.090 | 1.000 | 0.552 | 0.714 | 0.994 |
| | `hap-ibd` | 0.025 | 0.089 | 1.000 | 0.544 | 0.712 | 0.995 |
| | `iLASH` | 0.032 | 0.245 | 0.998 | 0.445 | 0.527 | 0.997 |
| | `RaPID` | 0.977 | 0.986 | 0.913 | 0.998 | 0.999 | 0.079 |
| | `TPBWT` | 0.995 | 0.991 | 0.316 | 0.984 | 0.999 | 0.845 |

**Running time and memory.** Table A2 gives the cost of each method without genotyping error. The linear methods ran on the node of Section 5.1, and GIBD on an AMD EPYC 75F3 node with 512 GB of memory. GIBD is slower because it builds the GBWT and reports every pairwise-maximal match, while the linear methods work on a matrix of biallelic sites.

Table A2: Wall-clock time and peak memory in the experiments of the simulation of Tang et al. without genotyping error. For GIBD, building the GBWT and the search are given separately.

| | All sites | | MAF $\geq 0.1$ | |
|---|---|---|---|---|
| Method | Time (s) | Memory (GB) | Time (s) | Memory (GB) |
| GIBD (GBWT build / search) | 167 / 743 | 10.3 / 18.9 | 35 / 285 | 2.2 / 2.9 |
| `hap-ibd` | 41 | 0.7 | 9 | 0.6 |
| `iLASH` | 50 | 1.6 | 8 | 0.3 |
| `RaPID` | 39 | 0.02 | 85 | 0.06 |
| `TPBWT` | 284 | 0.1 | 18 | 0.1 |

### C Simulated haplotypes from HPRC assemblies

This simulation underlies Section 5.4 on T2T-CHM13 and Section 5.5 on GRCh38, with the same rules, pipeline and quality control on both references. The haplotypes carry the insertions and deletions of real HPRC v2 assemblies and planted inversions, so the IBD segments and their SVs are known.

**Region and founders.** We used chr22:21–38 Mb on GRCh38 (21.7–38 Mb on T2T-CHM13), the longest interval of chromosome 22 in which at least 400 haplotypes of the HPRC v2 Minigraph-Cactus graph [2, 3] are each assembled as a single contiguous walk. It spans 29.9 cM on the T2T-CHM13 map [4] and 31.9 cM on the HapMap GRCh38 map [5], and it contains the 22q11.2 segmental duplications. Of the 418 contiguous walks on T2T-CHM13 and 416 on GRCh38, 27 carry a somatic V–J deletion at the IGL locus from the lymphoblastoid cell lines of HPRC [6], and we excluded them. Each replicate drew 200 founders from the remaining 391 and 389 walks.

**Ancestry.** We simulated ancestry with `msprime` 1.2.0 [7] under the discrete-time Wright–Fisher model [8], with the population and sample of Section 5.4 and recombination from the genetic map of the reference. Each lineage alive 40 generations ago was assigned a distinct founder walk. A sample haplotype joined the founder sub-walks it inherited at the first reference node after each breakpoint that both founder walks traverse forward.

**New variation.** Point mutations arose at $1.25 \times 10^{-8}$ per bp per generation in the last 40 generations, with the new base uniform among the other three [1]. All carriers of a mutation shared one copy of the node with the base changed. Each replicate also had 20 planted inversions of 5–100 kb (log-uniform), at least 500 kb apart and from the ends of the region. Each inversion was placed on a branch of the local tree with at least two sample descendants, chosen with probability proportional to the branch length. We kept an inversion only if the local tree was the same across it and at least two haplotypes of the same founder were not under the branch, so that carriers and non-carriers shared IBD across it. In each carrier, the sub-walk across the inversion was reversed. The condition with assembly error added independent substitution errors at $10^{-5}$ per bp per haplotype (QV50), more than reported for HPRC assemblies [9], on the same ancestry, mutations and inversions.

**Truth.** Known IBD segments are the `tskit` IBD segments between sample haplotypes whose most recent common ancestor is younger than 40 generations, merged while the ancestor is unchanged and across gaps shorter than 5 kb, following Tang et al. [1]. The SVs of a known segment are those of its shared founder walk, called by GIBD's rule. Between consecutive reference nodes, a haplotype carries an insertion if its bases exceed the reference distance by at least 50 bp and a deletion if they fall short by at least 50 bp. The planted inversions and their carriers complete the SV truth.

**Quality control.** Before any method ran, each replicate had to pass checks that do not depend on the methods. Every walk uses existing edges. For 20 random haplotypes, the spelled sequence equals an independent reconstruction from the founder sequences, breakpoints, mutations, errors and inversions. For 1,000 random known segments, both haplotypes copy the same founder. At least 99% of the planted point mutations outside the inversions appear in the VCF with their exact carriers.

**Linear methods.** On GRCh38, we also ran the linear methods on the VCF that the consortium's Minigraph-Cactus pipeline [3] derives from the simulated graph. The pipeline runs `vg deconstruct` (vg 1.74.1) [10], `vcfbub` (0.1.0) to remove sites of more than 100 kb, `vcfwave` (vcflib 1.0.15) [11] to decompose complex alleles, and `bcftools` 1.24 [12] to normalize and split records into biallelic ones. At a position with several records, we kept the one with the largest minor allele count, since `hap-ibd` and `IBD_BM` need one marker per position. These markers also gave GIBD's genetic map, so all methods measured length in the same coordinates. `hap-ibd` [13] ran with its default parameters, and `RaPID` [14], `iLASH` [15] and `TPBWT` [16] with the parameters of Tang et al. [1], all with the 2 cM output threshold.

**Measures.** For each planted inversion $[a, b]$, we took the haplotype pairs whose known segment covers $[a - 50\,\text{kb},\ b + 50\,\text{kb}]$. A pair differs at the inversion if exactly one haplotype carries it, and the other pairs form the control. GIBD spans a pair if one reported segment covers the same interval. For pairs that differ, we report the fraction spanned and the fraction of spanning segments labeled as differing by the inversion. For control pairs, we report how many GIBD labeled as differing. For pairs of carriers, Detected is the fraction labeled with the shared inversion. For both inversion rows, precision is the fraction of inversion labels that are correct. Insertion and deletion labels are evaluated on matched segments, where a reported and a known segment each overlap at least half of the other. We ran five replicates per condition.

**Results.** Table A3 gives the SV labels of GIBD on GRCh38. GIBD detected 72% of the known segments with an insertion or deletion without assembly error and 63% with it. When exactly one haplotype carries an inversion, GIBD spanned 61% of the pairs, and it spanned 63% of the about 119,700 control pairs. When both haplotypes carry the inversion, GIBD labeled it as shared for 80% of the pairs. GIBD's power on known segments that carry an SV was 0.618 and 0.532 at 2–3 cM and 0.893 and 0.802 at $\geq 7$ cM, without and with assembly error. Table A4 gives the power and accuracy of the linear methods on the same simulation, on all known segments rather than only those that carry an SV.

Table A3: SV labels of GIBD in the simulation, without / with assembly error (five replicates each). Accuracy is that of 2–3 cM segments with the label.

| Label | Precision | Recall | Detected | Accuracy [2,3) cM |
|---|---|---|---|---|
| Insertion | 1.000 / 1.000 | 1.000 / 1.000 | 0.724 / 0.633 | 0.996 / 0.996 |
| Deletion | 1.000 / 1.000 | 1.000 / 1.000 | 0.725 / 0.636 | 0.995 / 0.996 |
| Inversion, one haplotype carries it | 1.000 / 1.000 | 1.000 / 1.000 | 0.613 / 0.531 | 1.000 / 1.000 |
| Inversion, both carry it | 1.000 / 1.000 | – | 0.799 / 0.784 | 1.000 / 1.000 |

Table A4: Power (P) and accuracy (A) of the linear methods from `IBD_BM` by known segment length (cM) in the GRCh38 simulation, on all known segments, mean over five replicates per condition.

| | Without error | | | With assembly error | | |
|---|---|---|---|---|---|---|
| Method | P [2,3) | P $\geq 7$ | A [2,3) | P [2,3) | P $\geq 7$ | A [2,3) |
| `hap-ibd` | 0.616 | 0.890 | 0.996 | 0.524 | 0.804 | 0.997 |
| `iLASH` | 0.557 | 0.868 | 0.997 | 0.013 | 0.160 | 0.995 |
| `RaPID` | 0.996 | 0.998 | 0.990 | 0.991 | 0.995 | 0.993 |
| `TPBWT` | 1.000 | 1.000 | 0.992 | 0.983 | 0.968 | 0.992 |

**Limitations.** The simulation reuses the real HPRC graph, so it does not model graph-construction error, which affects all methods alike. It covers one founder population and one region. The planted inversions lack the flanking inverted repeats of many real inversions, and assembly errors are substitutions only.

### D Platinum Pedigree

**Data and graphs.** We used the phased assemblies of the 23 openly released members of CEPH 1463 (46 haplotypes) [17, 18]. For chromosomes 17 and 20, we assigned contigs to the chromosome by their alignment to GRCh38 [19] and built a pangenome graph with the Minigraph-Cactus recipe of the HPRC, with GRCh38 as the reference [3].

**Inheritance truth.** We inferred the inheritance of each child haplotype with the Lander–Green hidden Markov model [20] over the parents' heterozygous SNPs in the consortium's phased family truthset [18]. The truthset covers NA12877, NA12878 and five of their children. On chromosome 20, the model places 11 crossovers in 10 meioses, each within 33 kb, and defines 55 known IBD segments between parents and children and between siblings (54 of at least 2 cM). On chromosome 17, it places 15 crossovers and defines 64 known segments. Members with no common ancestor in the pedigree share no IBD.

**Input of GIBD.** On chromosome 20, inherited haplotypes differ at 24,706 sites over 21 parent–offspring blocks of 621 Mb in total (40 per Mb). Of these, 82% are indels of 1–49 bp, and 96% of those change the length of a homopolymer or a short tandem repeat, the residual error of HiFi assembly [21]. On the unmodified graph, GIBD found almost no IBD (power 0.007 at seed and extension lengths of 2 and 1 cM). As is conventional for IBD detection, GIBD therefore read only SNPs and SVs. In its graph, each haplotype follows the reference allele wherever it differs from it by an indel of 1–49 bp or by adjacent substitutions. SNPs and alleles of at least 50 bp are kept, so the SV labels are unchanged.

**Parental haplotype at SVs.** At a callset SV where a parent is heterozygous, the child's allele shows which parental haplotype the child inherited. For each child haplotype and each such SV inside a GIBD parent–offspring segment, we checked that the child carries the allele of the parental haplotype in the segment. On chromosome 20, there are 5,030 such pairs of a child haplotype and an SV, and GIBD's segments cover 2,601, 3,235 and 3,761 of them at the three seed and extension lengths of Table 4.

**SV labels against the callset.** GIBD's labels for a segment are the SVs that both haplotypes carry. We compared them with the consortium's SV callset [18], which merges read-based and assembly-based calls, using Truvari [22] with reference distance 500 bp and size similarity 0.7. Recall counts the callset SVs that both haplotypes carry inside known segments that GIBD covered. At seed and extension lengths of 2 and 1 cM, lenient matching (2,000 bp, 0.3) raised precision only to 0.63 for insertions and 0.76 for deletions. Both haplotypes of a known segment carry the same callset allele at all 11,256 records inside it. An SV lies in a tandem repeat if it overlaps the UCSC simpleRepeat track (Tandem Repeats Finder [23]) within 50 bp. Outside tandem repeats, for callset SVs that its assembly-based callers also found, the graph's recall on single haplotypes is 0.84 for insertions and 0.92 for deletions. The two haplotypes of a known segment were assembled independently. Inside known segments, 93–94% of the graph's SVs occur identically on both haplotypes, including 90% of those absent from the callset.

**Chromosome 17 and the 17q21.31 inversion.** Table A5 gives the results on chromosome 17. A haplotype carries H2 if one of its contigs aligns to both flanks (45.0–45.45 and 46.45–46.9 Mb) in one orientation and to the core (45.775–46.125 Mb) in the other, and H1 if all three align in the same orientation. This called 9 haplotypes H2 and 30 H1, and left 7 without a call. The inheritance truthset covers only seven members, so we evaluated every GIBD segment that spans chr17:45.6–46.3 Mb between relatives in the published pedigree. A segment between two H2 haplotypes should carry the inversion label, and a segment between two H1 haplotypes the reference allele. No segment joined an H1 and an H2 haplotype.

Table A5: GIBD in the Platinum Pedigree, chromosome 17, on known segments of at least 2 cM. The last row counts the GIBD segments across 17q21.31 between two haplotypes called H1 or H2 that were labeled as expected, with wrong labels in parentheses. The remaining segment at 2/1 and at 0.5/0.25 has an end inside 17q21.31 and carries no label.

| Seed / extension length (cM) | 2 / 1 | 1 / 0.5 | 0.5 / 0.25 |
|---|---|---|---|
| Power | 0.358 | 0.538 | 0.632 |
| Segments between unrelated members | 0 | 0 | 0 |
| SVs on the correct parental haplotype | 1,747 / 1,747 | 2,614 / 2,614 | 3,163 / 3,163 |
| 17q21.31 segments labeled as expected (wrongly) | 4 / 5 (0) | 23 / 23 (0) | 44 / 45 (0) |

### E Pseudocode of GIBD

Algorithm A1 enumerates the gMEMs (Sections 3.2 and 3.3), and Algorithm A2 turns them into SV-labeled IBD segments (Section 3.4).

**Algorithm A1** Enumeration of gMEMs

**Require:** GBWT of the indexed paths $\mathcal{P}^{\pm}$, minimum length $b_m$ in bases
**Ensure:** gMEMs between distinct haplotypes, of length at least $b_m$ (lower for gMEMs traversed in opposite orientations)

**for** each oriented node $v \in V^{\pm}$ **do**
  **if** two rows of $\mathsf{BWT}_v$ store different successors or both store \$ **then** ▷ $v$ is right-maximal
    locate the rows of $\mathsf{BWT}_v$ with FastLocate
    Extend($(v)$, $[0, |\mathsf{BWT}_v| - 1]$)
  **end if**
**end for**

**procedure** Extend($Q$, $[s, e]$) ▷ $[s, e]$: rows of $\mathsf{BWT}_v$ of the occurrences of $Q$
  **if** two occurrences of $Q$ have different preceding nodes or both start a path **then** ▷ left-maximal
    Report($Q$, $[s, e]$)
  **end if**
  **for** each node $u$ preceding an occurrence of $Q$ **do**
    $[s_u, e_u] \leftarrow$ rows of the occurrences of $uQ$, by a successor lookup in $\bar{Q}$
    **if** two rows of $[s_u, e_u]$ store different successors or both store \$ **then**
      Extend($uQ$, $[s_u, e_u]$)
    **end if**
  **end for**
**end procedure**

**procedure** Report($Q$, $[s, e]$)
  group the rows of $[s, e]$ by the preceding node $u$ and the successor $w$ of their occurrences
  **for** each pair of groups $(u, w)$ and $(u', w')$ that satisfy Equation (1) **do**
    **for** each pair of distinct rows $k$ of group $(u, w)$ and $k'$ of group $(u', w')$ **do**
      $(P, i), (P', i') \leftarrow$ occurrences of $Q$ at rows $k$ and $k'$, from the located rows
      **if** $P$ and $P'$ belong to distinct haplotypes and $Q$ has at least $b_m$ bases, or at least a lower floor if $P$ and $P'$ traverse $Q$ in opposite orientations **then**
        output the gMEM $\{(P, i), (P', i')\}$ if the lower-numbered haplotype is read forward (Lemma A1) and no gMEM of $Q$ on the same two haplotypes with the same relative orientation was output
      **end if**
    **end for**
  **end for**
**end procedure**

**Algorithm A2** SV-labeled IBD segments

**Require:** haplotype paths $\mathcal{P}$, the path $R$ of the reference genome, genetic map, minimum number of haplotypes $m$ per node, minimum gMEM length $b_m$ in bases, seed length $s$ and extension length $x$ in cM, maximum gap $d$ in bases, minimum IBD length $c$ in cM, minimum SV length $b_v$ in bases, length tolerances $f$ and $f_{bp}$

**Ensure:** IBD and HBD segments, each labeled with the insertions, deletions and inversions that both haplotypes carry

**for** each path $P \in \mathcal{P}$ **do** ▷ mask rare alleles
    replace each node traversed by fewer than $m$ haplotypes with the subpath that most haplotypes traverse between the nearest preceding and succeeding nodes traversed by at least $m$ haplotypes
**end for**
build the GBWT of the masked paths and their reverse complements
$\mathcal{M} \leftarrow$ gMEMs of the masked paths (Algorithm A1)
**for** each pair of haplotypes $(h, h')$ with gMEMs in $\mathcal{M}$ **do**
    place each gMEM of $(h, h')$ on $R$ using shared nodes, with its length in cM from the genetic map
    join the inverted gMEMs between two consecutive gMEMs of $(h, h')$ to them if there are at most two and every gap is at most $d$ bases, and mark the joined region inverted
    join consecutive gMEMs of at least $x$ that are separated by at most $d$ bases into candidate segments
    **for** each candidate segment that contains a gMEM of at least $s$ and spans at least $c$ **do**
        label the segment with every SV of at least $b_v$ bases that $h$ and $h'$ both carry: same type, overlapping intervals on $R$, and lengths within $\max\{f_{bp}, f \cdot \text{longer length}\}$; label it inverted if it contains an inverted region
        report the segment, as HBD if $h$ and $h'$ are the two haplotypes of one sample and as IBD otherwise
    **end for**
**end for**

### References

[1] Kecong Tang, Ardalan Naseri, Yuan Wei, Shaojie Zhang, and Degui Zhi. Open-source benchmarking of IBD segment detection methods for biobank-scale cohorts. *GigaScience*, 11:giac111, 2022.

[2] Human Pangenome Reference Consortium. HPRC pangenome resources, second release: Minigraph-Cactus graphs and assemblies. `https://github.com/human-pangenomics/hpp_pangenome_resources`, 2024. Data release, accessed 2026.

[3] Glenn Hickey, Jean Monlong, Jana Ebler, Adam M Novak, Jordan M Eizenga, Yan Gao, Human Pangenome Reference Consortium, Haley J Abel, Lucinda L Antonacci-Fulton, Mobin Asri, et al. Pangenome graph construction from genome alignments with Minigraph-Cactus. *Nature Biotechnology*, 42(4):663–673, 2024.

[4] Joseph L. Lalli, Andrew N. Bortvin, Rajiv C. McCoy, and Donna M. Werling. A T2T-CHM13 recombination map and globally diverse haplotype reference panel improves phasing and imputation. *bioRxiv*, 2025.
